# COVID-3D: An online resource to explore the structural distribution of genetic variation in SARS-CoV-2 and its implication on therapeutic development

**DOI:** 10.1101/2020.05.29.124610

**Authors:** Stephanie Portelli, Moshe Olshansky, Carlos H.M. Rodrigues, Elston N. D’Souza, Yoochan Myung, Michael Silk, Azadeh Alavi, Douglas E.V. Pires, David B. Ascher

## Abstract

The emergence of the COVID-19 pandemic has spurred a global rush to uncover basic biological mechanisms, to inform effective vaccine and drug development. Despite viral novelty, global sequencing efforts have already identified genomic variation across isolates. To enable easy exploration and spatial visualization of the potential implications of SARS-CoV-2 mutations on infection, host immunity and drug development we have developed COVID-3D (http://biosig.unimelb.edu.au/covid3d/).

## MAIN MANUSCRIPT

Declared a global pandemic on March 11^th^ 2020^1^, COVID-19 has become the most recent modern-day global health challenge, infecting almost 5 million people and claiming over 300,000 lives to date^2^. Consequently, the scale of its humanitarian and economic impact has driven academic and pharmaceutical efforts to develop vaccines and antiviral treatments. Current efforts include 118 active vaccine candidates^3^, and numerous more endeavours to identify biologics and small molecule treatments.

One further challenge in controlling COVID-19 is the accumulation of variation across genes. Sources indicate that SARS-CoV-2 is mutating at about 2 variants/month^4^, but the potential implications of these on molecular diagnostics and the development of candidate vaccines and treatments remain poorly explored. Fortunately, the continuous exponential increase in the amount of SARS-CoV-2 genome sequence data and structural information available provides the opportunity to analyse both data sources concomitantly. This provides a unique opportunity to not only understand how variants might affect patient outcomes, but also anticipate and minimise their potential role in viral escape through early incorporation within the development pipeline.

To facilitate this, we have developed a comprehensive online resource, COVID-3D, to enable analysis and interpretation of variants detected in over 45,000 SARS-COV-2 genomic sequences^5^ (Figure S1). We have mapped these circulating variants to their respective protein sequences and structures of the SARS-COV-2 proteins derived from available experimental information, permitting a direct comparison of variant clustering between the two representations. Our interactive 3-D viewer enables fast and intuitive spatial visualisation of SARS-CoV-2 variants, highlighting their potential impacts on protein structure and interactions^6–12^ (Figure S2-S5). This is particularly useful for analysing sites being currently targeted by potential therapeutics. To further enhance therapeutic discovery efforts, we have included drug binding potential^13,14^ and predicted antigenicity maps^15,16^ of the structures, which permit rational selection of target sites and compound design specifically avoiding already circulating variants (Figure S3). Finally, combining this structural information with evolutionary and population variation analysis can further help identify sites less likely to accommodate mutations in the future. To illustrate this, COVID-3D was used to provide insights into the two main therapeutic targets- the Spike protein and Main Proteinase.

The SARS-CoV-2 spike protein binds to human Angiotensin-converting enzyme 2 (ACE2) mediating cell entry. Subsequently, the ACE2 receptor binding domain has been the main target of most vaccine programs. Measures of selective pressure suggest that the spike protein is one of the viral proteins most tolerant to the introduction of mutations^17,18^ (Table S1). Upon closer inspection (http://biosig.unimelb.edu.au/covid3d/protein/QHD43416/CLOSED), it is evident that despite SARS-CoV-2 only being discovered 6 months ago, we can already see significant variation across the protein surface, including in predicted epitope regions in the receptor binding domain (Figure 1B). Of these variants, the D614G mutation is present in two-thirds of the sequenced strains, although its actual significance remains unclear despite initial suggestions at increasing transmissibility^19^. The residue is located far from the ACE2 interface (73 Å), and was predicted to have a mildly stabilising (DUET^8^ 0.5 kcal/mol; SDM^7^ 2.3 kcal/mol) effect on protein stability, and hence a minimal fitness cost^20^. It was, however, predicted to alter protein dynamics and the interactions between the subunits (4.4 Å from the interface. mCSM-PPI2^11^ −0.5 Kcal/mol for the closed form versus −0.35 Kcal/mol for the open form), which could affect the equilibrium between open and closed states.

**Figure 1.**
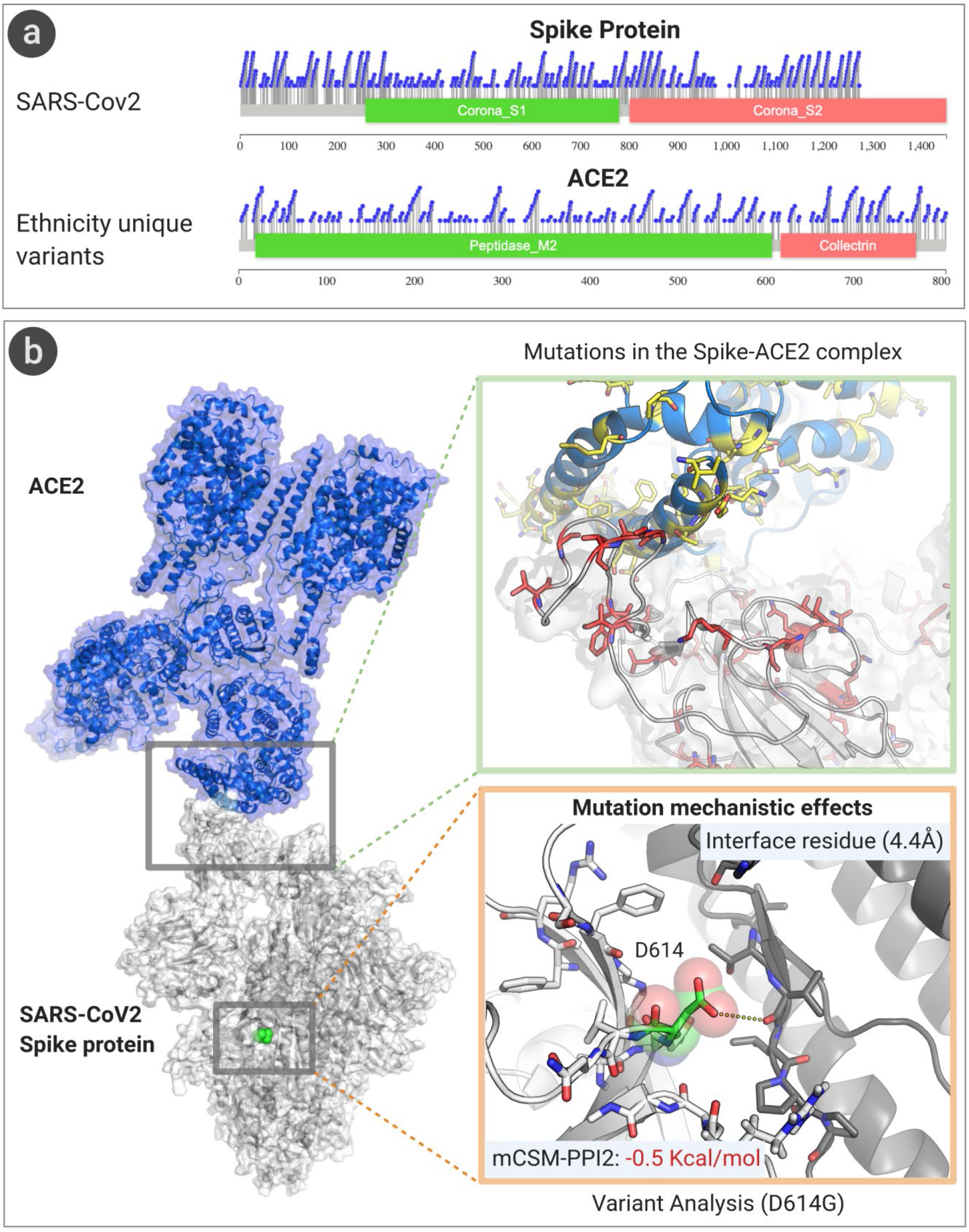
Population variation across the Spike-ACE2 complex. a) Lollipop plots of circulating missense variants in the SARS-CoV2 Spike protein and ethnically unique missense variants in human ACE2 illustrate the broad spread of changes across the proteins. b) When they are visualised spatially, there are a number of variants seen at the ACE2-Spike interface that are predicted to impact on the binding affinity. One of the most prevalent circulating SARS-CoV2 Spike variants, D614G, is located far from the ACE2 interface, but close to the Spike trimer interface and is predicted to lead to structural perturbations.

Interestingly, when we look at population specific variants across ACE2, we see a number of ethnic group specific variants across the interface recognised by Spike (Figure 1A). Evaluation of their consequences using mCSM-PPI2^11^, which has been experimentally validated on this protein system^21^, reveals potential significant effects on the binding affinity of Spike, opening up further work to explore how this influences the severity and progression of COVID-19.

Apart from Spike, the Main Proteinase (http://biosig.unimelb.edu.au/covid3d/protein/QHD43415_5/APO) has also attracted a lot of therapeutic development efforts, as a target for small molecule development. The Main Proteinase, however, is not particularly intolerant to missense variants (Table S1), which may promote the emergence of resistant variants. The structures show that there are already a number of circulating variants present in the drug binding site that could have implications on efficacy (Figure 2A). Using COVID-3D, we have leveraged the wealth of SARS-CoV-2 genomic sequences to calculate measures of mutational tolerance, which revealed a number of proteins under strong purifying selection (Table S1). This includes the Helicase, NSP7, NSP8, NSP9 and ExoN, which may make novel attractive drug targets with few circulating variants seen near the druggable pockets (Figure 2B).

**Figure 2.**
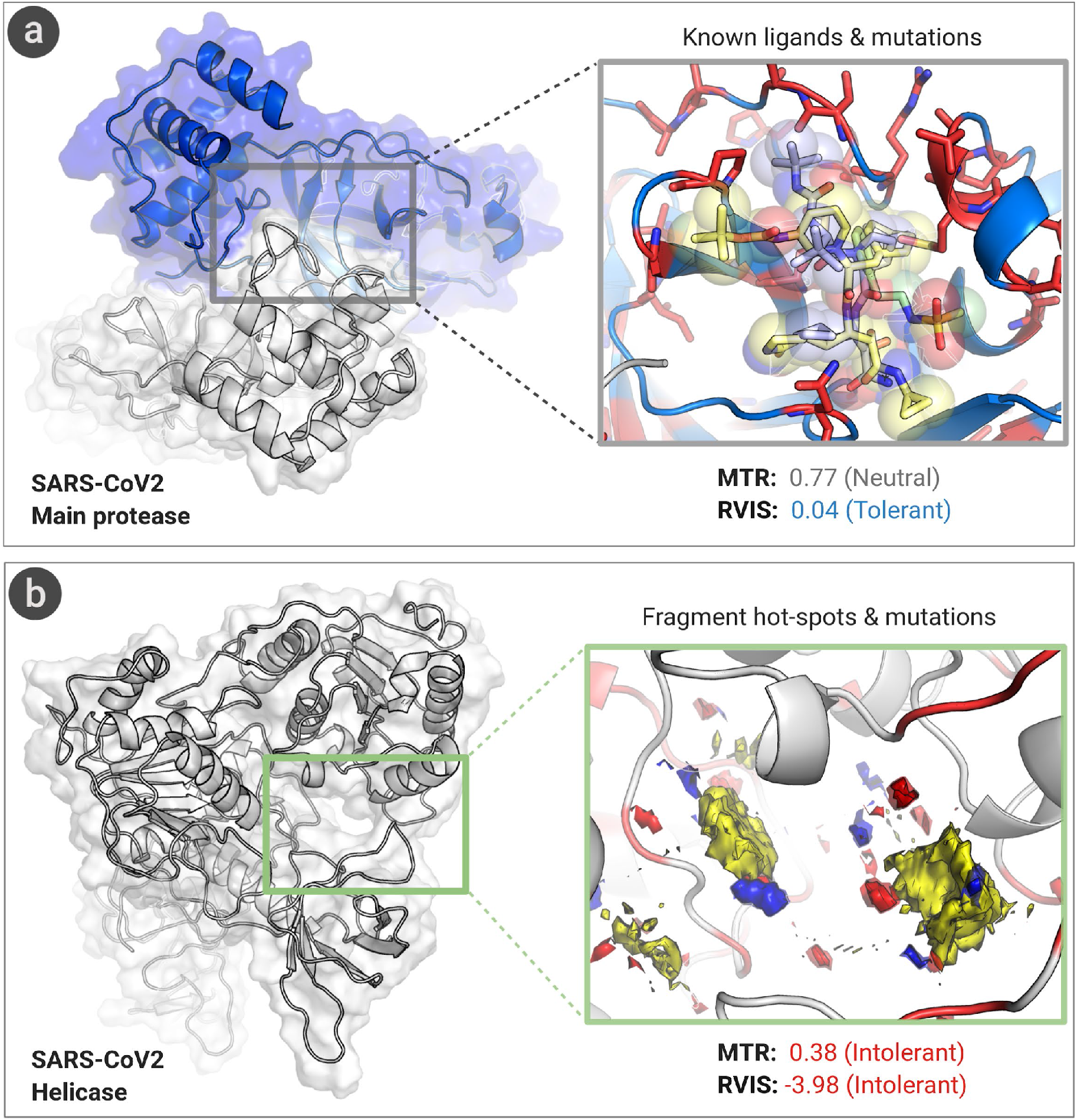
Visualisation of circulating variants relative to druggable pockets. a) The Main Proteinase is neutral to the introduction of missense variants. Circulating variants (red sticks) have already been seen in close proximity to known inhibitors, and are likely to affect binding. This suggests that resistant mutations could be selected for with widespread use. b) The Helicase is one of the genes most intolerant to missense variation. Mapping the fragment binding potential reveals pockets with apolar (yellow), hydrogen bond donor (blue), and hydrogen bond acceptor (red) potential. While some variation has been observed close by, optimisation of interactions to avoid these sites could reduce the potential for future resistance.

COVID-3D provides an easy to use bridge between genomic information and structural insight to better guide our biological understanding and treatment efforts. As new structural and sequence data becomes available, COVID-3D will be periodically updated to enable their integration into ongoing efforts to understand and combat SARS-CoV-2.

## METHODS

### Mapping genetic variants

High quality SARS-CoV-2 genomic sequences were obtained from GSAID^5^ and the COG-UK Consortium. Sequences were aligned using blastn to the reference genome (NC_045512.2), and synonymous and missense variants for each mature protein curated.

Human ACE2 and B0AT1 population variants were obtained from gnomAD^22^, UK10K project^23^, KRGDB^24^, 4.7KJPN Tohoku Medical Megabank Project^25^, and the UK BioBank^26^. Population specific variants were jointly called using PLINK and BCFTools, and all variants were converted to GRCh38/Hg38 genomic coordinate positions. Ensembl’s VEP (version 97) was used to identify missense and synonymous variants.

### Gene-level Essentiality Scores for SARS-CoV-2 proteins

A per-gene MTR score^17^ was calculated for each of the 25 SARS-CoV-2 CDS sequences (15 mature peptides derived from the polyprotein and 10 additional proteins). Observed variation was collapsed to unique missense and synonymous observations. The proportion of missense variants was compared with the expected proportion under neutrality from all possible variants from the NCBI reference CDS sequence.

Another metric, RVIS, was used to provide an alternate report of the essentiality of each gene^18^. Common functional variants (with Minor Allele Frequency of 0.01% or greater) were tallied for each gene. The number of common functional variants within each gene were regressed onto the number of all variants observed in that gene regardless of frequency using simple linear regression. The studentized residuals were extracted to calculate the RVIS score.

### Structural modelling

All protein fasta sequences within the SARS-CoV-2 genome were obtained from GenBank (MN908947.3) and blast against the RCSB protein data bank^27^ to identify experimental crystallographic SARS-CoV-2 structures or suitable templates for homology modelling. Experimental crystal structures were saved as biological assemblies, and optimized in Maestro (Schrodinger suite, v. 2017-4). Homology models were generated using Modeller^28^ and I-TASSER^29^ and optimized using Maestro. Structures were validated using Meastro Protein Preparation Wizard and Molprobity.

### Structural characterisation

Potential linear and structural epitopes predicted using DiscoTope 2.0^15^ and ElliPro^16^ respectively, pockets detected using GHECOM^14^, and fragment-binding hot-spot potentials using CCDC^13^. Surface electrostatics partial charges were generated using CHARMM^30^. Normal Mode Analysis was performed for each protein using DynaMut^10^ and Molecular Dynamic simulations using Discovery Studio. All intra- and inter-molecular interactions of missense variants were calculated using Arpeggio^6^. The molecular consequences of variants on protein stability were assessed using mCSM-Stability^9^, SDM^7^ and DUET^8^, and on protein-protein interactions using mCSM-PPI2^11^. Changes in interaction affinities to ligands and nucleic acids were calculated using mCSM-lig^12^ and mCSM-DNA^9^ where applicable. The MTR score^17^ for ACE2 and BOAT1 was calculated for each protein position with a sliding window of 41-codon for every ethnic population MTR scores and mapped onto the ACE2-B0AT1-Spike structure

### COVID-3D web interface

We have implemented COVID-3D as a user-friendly and freely available web server (http://biosig.unimelb.edu.au/covid3d/). The Materializecss framework version 1.0.0 was used to develop the server front end, while the back-end was built in Python using the Flask framework version 1.0.2. The server is hosted on a Linux server running Apache 2.

## ACKNOWLEDGEMENTS

S.P, C.H.M.R, and Y.M. were supported by the Melbourne Research Scholarship. D.B.A and D.E.V.P were funded by a Newton Fund RCUK-CONFAP Grant awarded by The Medical Research Council (MRC) and Fundação de Amparo à Pesquisa do Estado de Minas Gerais (FAPEMIG) [MR/M026302/1] and Conselho Nacional de Desenvolvimento Científico e Tecnológico (CNPq); the Jack Brockhoff Foundation [JBF 4186, 2016]; Wellcome Trust (200814/Z/16/Z), and an Investigator Grant from the National Health and Medical Research Council (NHMRC) of Australia [GNT1174405]. Supported in part by the Victorian Government’s OIS Program. This research has been conducted using the UK Biobank Resource under Application Number 50000.

## AUTHOR CONTRIBUTIONS

S.P. was responsible for structure curation, homology modelling, and structural characterisation. M.O. was responsible for curating SARS-CoV-2 variants. C.H.M.R. was responsible for developing the website. Y.M. performed the molecular dynamics and assisted with the website. E.D. was responsible for curating the human population variants. E.D. and M.S. were responsible for calculating intolerance scores. A.A. assisted with SARS-CoV-2 genomic curation. D.E.V.P. was responsible project supervision and for Spike protein characterisation. D.B.A. designed and supervised all aspects of the project. All authors assisted with manuscript writing.

## ADDITIONAL INFORMATION

Supplementary Information is available.

