## Supplementary Information for "COVID-3D: An online resource to explore the structural distribution of genetic variation in SARS-CoV-2 and its implication on therapeutic development"

**Table S1.** Using available genetic information to measure mutational tolerance of SARS-CoV-2 proteins.

| Protein | Uniprot | Missense Tolerance Ratio | | Residual Variation Intolerance Score |
|---|---|---|---|---|
| | | Score | Adjusted P-value | |
| Non-structural protein 9 (NSP9) | P0DTD1 | 0.336 | 8.36E-19 | -0.704 |
| Helicase (Hel) | P0DTD1 | 0.375 | 3.72E-80 | -3.980 |
| Guanine-N7 methyltransferase (ExoN) / Proofreading exoribonuclease | P0DTD1 | 0.384 | 5.30E-74 | -1.999 |
| Non-structural protein 7 (NSP7) | P0DTD1 | 0.419 | 6.35E-06 | -0.237 |
| Non-structural protein 8 (NSP8) | P0DTD1 | 0.420 | 1.80E-20 | -0.661 |
| Non-structural protein 10 (NSP10) | P0DTD1 | 0.530 | 1.06E-04 | -0.243 |
| Envelope Small Membrane Protein | P0DTC4 | 0.637 | 2.59E-03 | -0.057 |
| Membrane Glycoprotein | P0DTC5 | 0.643 | 3.70E-06 | -0.167 |
| 2-O-methyltransferase (2-O-MT) | P0DTD1 | 0.643 | 1.09E-07 | -0.304 |
| Non-structural protein 4 (NSP4) | P0DTD1 | 0.669 | 1.01E-09 | -0.029 |
| Non-structural protein 6 (NSP6) | P0DTD1 | 0.684 | 1.40E-06 | -0.114 |
| Host translation inhibitor (NSP1) | P0DTD1 | 0.700 | 1.11E-04 | 0.060 |
| ORF6 | P0DTC6 | 0.718 | 2.96E-02 | 0.044 |
| RNA-directed RNA polymerase (RdRp) / Non-structural protein 12 (NSP12) | P0DTD1 | 0.733 | 9.28E-11 | -0.972 |
| ORF7b | (NOT CURATED) | 0.751 | 3.97E-02 | -0.071 |
| Main protease 3CL-PRO | P0DTD1 | 0.768 | 3.08E-04 | 0.037 |
| Papain-like protease (PLpro) | P0DTD1 | 0.779 | 1.65E-16 | 1.570 |
| Surface glycoprotein (Spike) | P0DTC2 | 0.781 | 9.87E-11 | 1.433 |
| ORF7a | P0DTC7 | 0.825 | 7.21E-03 | 0.615 |
| Uridylate-specific endoribonuclease (NendoU) | P0DTD1 | 0.833 | 4.56E-03 | -0.195 |
| Nucleocapsid protein | P0DTC9 | 0.838 | 4.80E-05 | 1.983 |
| Non-structural protein 2 (NSP2) | P0DTD1 | 0.876 | 1.45E-03 | 1.122 |
| ORF8 | P0DTC8 | 0.904 | 2.20E-01 | 0.467 |
| ORF3a | P0DTC3 | 0.960 | 3.74E-01 | 1.297 |
| ORF10 | A0A663DJA2 (NOT CURATED) | 1.008 | 1.00E+00 | 0.411 |

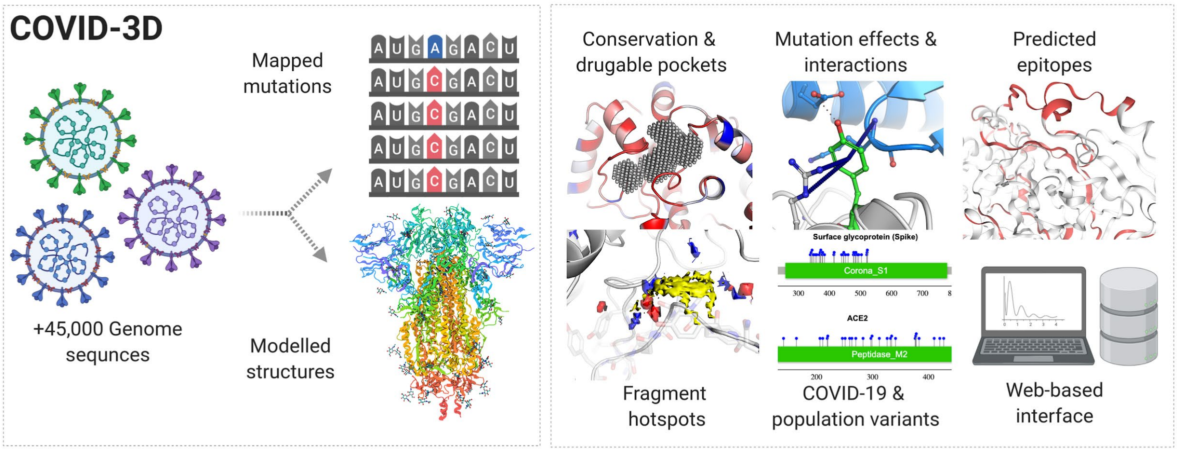


**Figure S1. The COVID-3D workflow.** SARS-CoV-2 genomes were curated from GISAID and COG, and mapped onto structures of the SARS-CoV-2 proteins. This enables their visualisation relative to druggable pockets and known interactions in order to guide understanding of their molecular consequences, and therapeutic development efforts.

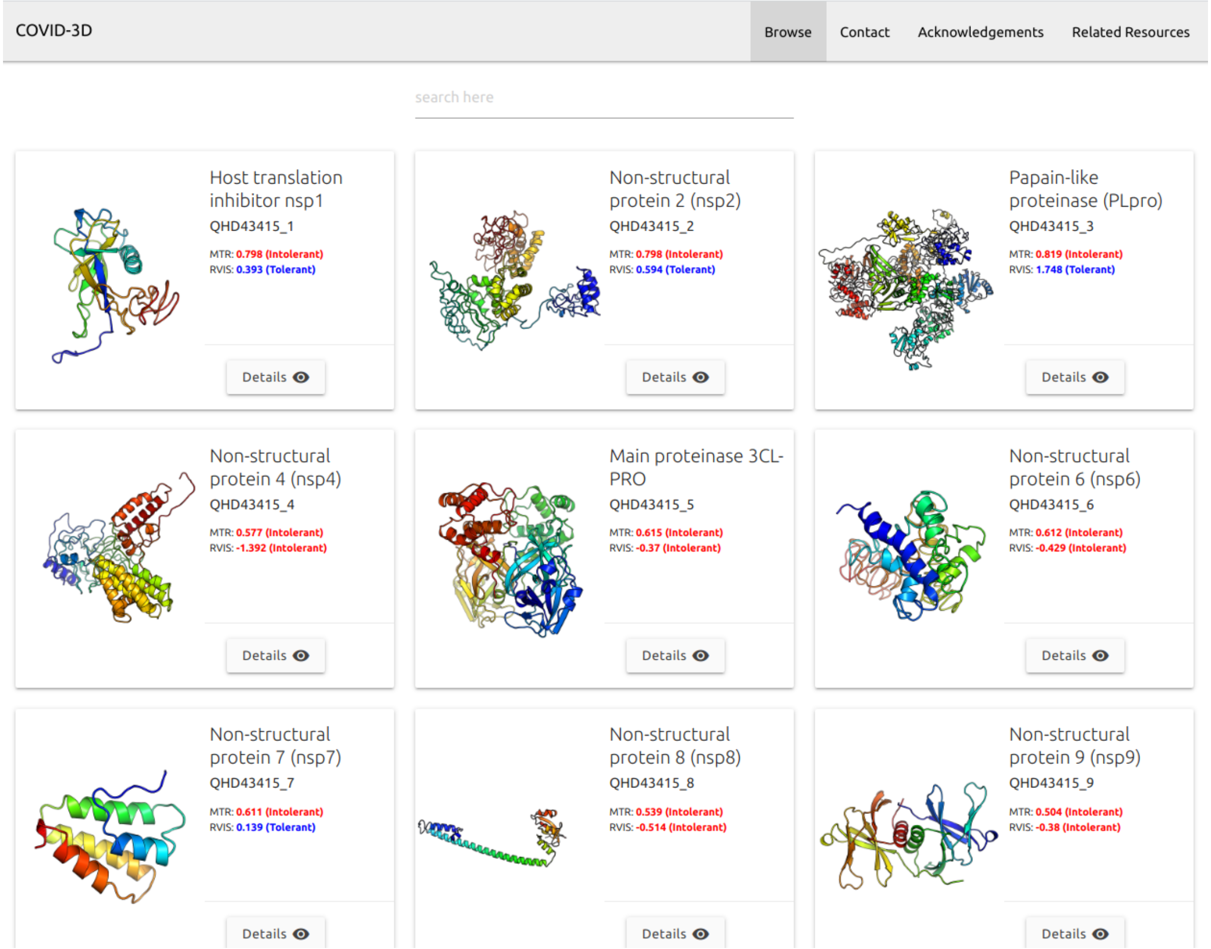


**Figure S2. COVID-3D listing page**. All proteins are displayed on separate cards containing name, uniprot accession number, missense tolerance ratio and residual variation intolerance scores. Structural characterisation and other analysis are available through a Detail button for each protein. A field on the top of the page allows for text search for any information on the card.

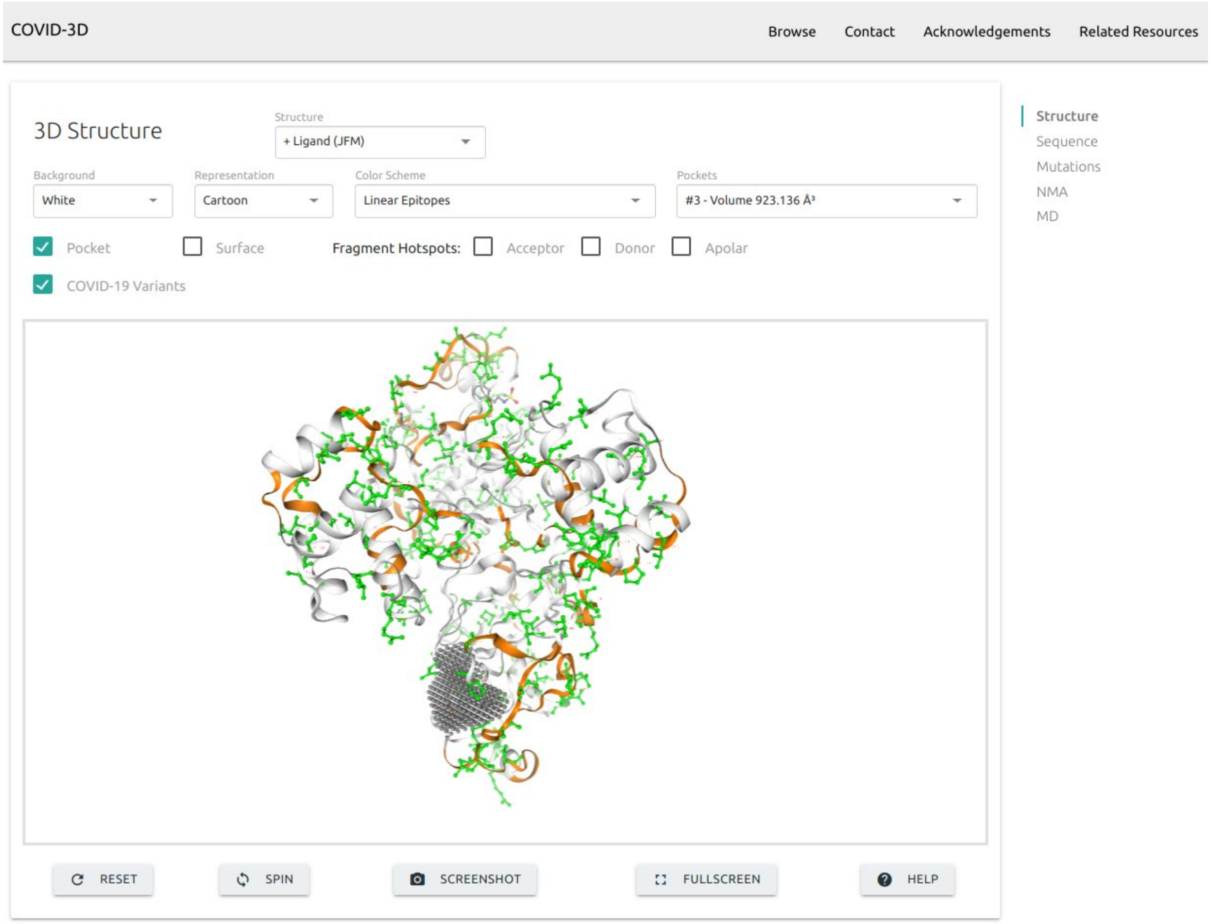


**Figure S3. Structure section of protein details page on COVID-3D**. Various controllers and switches are available for customising the interactive viewer. Here we see the structure of the Main proteinase 3CL-PRO (Uniprot: QHD43415_5) represented as cartoon with a pocket shown as little gray spheres grouped at the bottom, linear epitope regions coloured in orange and COVID-19 variants represented as balls and sticks and coloured in green.

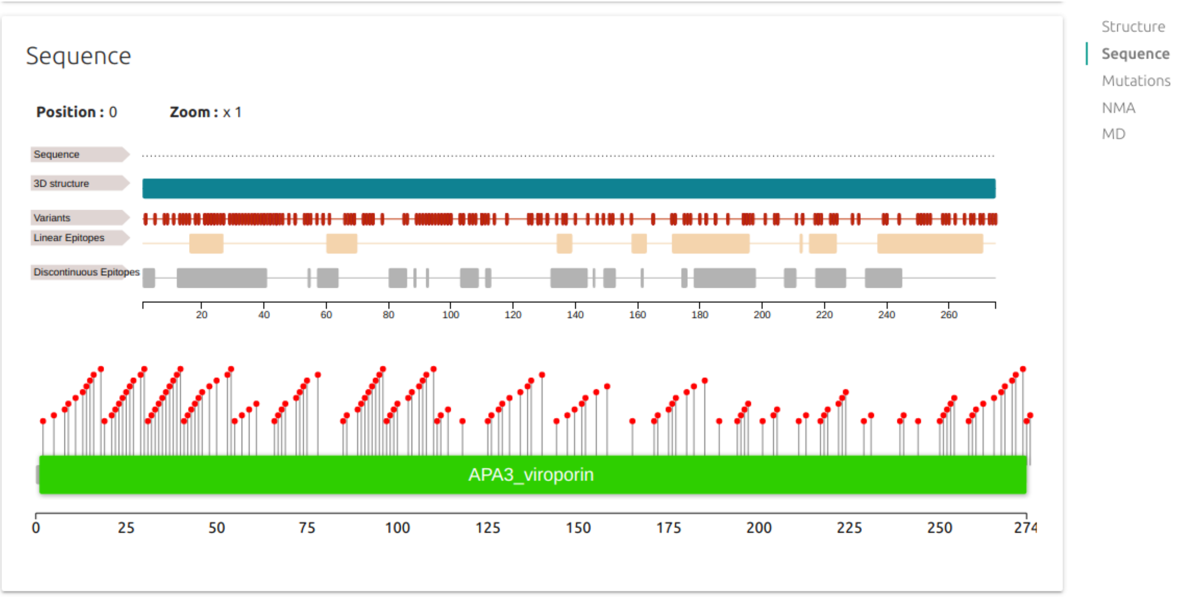


**Figure S4. Sequence section of protein details page on COVID-3D**. Protein sequence, regions mapped onto structure, variants and epitopes are combined on sequence viewer at the top of this section. If domain information is available a lollipop will be displayed at the bottom highlighting the domain region and the location for each variant (ORF3a - Uniprot: QHD43417).

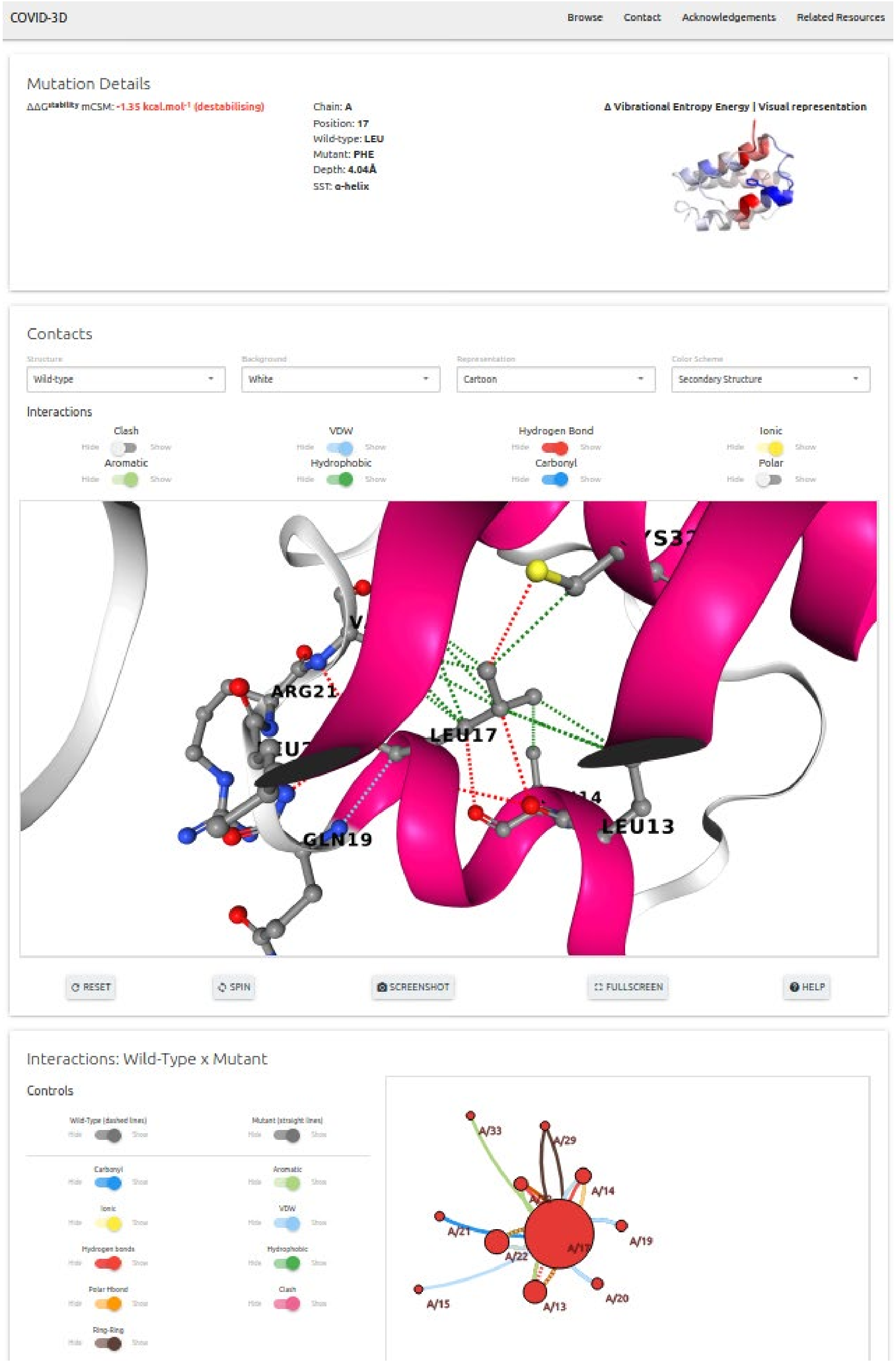


**Figure S5. Mutation details page on COVID-3D**. Stability, vibrational entropy changes and affinity changes (when applicable) caused by a particular variant are shown on the top panel alongside with information on the structural environment of the wild-type residue. An interactive 3D viewer where molecular contacts are displayed for wild-type residue is also available on the second panel. One can customise the structure using the controllers available and also select between wild-type and mutant structures. A 2D representation of the molecular contacts is also shown at the bottom of the page. Here we show the example of mutation L17P on chain A for the Non-structural protein 7 (nsp7 - Uniprot: QHD43415_7).
